# Novel antioxidant therapy with the immediate precursor to glutathione, γ-glutamylcysteine (GGC), ameliorates LPS-induced cellular stress in an *in vitro* cystic fibrosis model

**DOI:** 10.1101/2020.05.27.119990

**Authors:** Chris K Hewson, Alexander Capraro, Sharon L Wong, Elvis Pandzic, Bentotage S.M Fernando, Nikhil T Awatade, Ling Zhong, Gene Hart-Smith, Renee M Whan, Shane R Thomas, Adam Jaffe, Wallace J Bridge, Shafagh A Waters

## Abstract

**Introduction:** Glutathione deficiency and chronic bacterial inflammation exacerbates the oxidative stress damage to airways in cystic fibrosis. Improvements to current antioxidant therapeutic strategies are needed. Dietary supplement, γ-glutamylcysteine (GGC), the immediate precursor to glutathione, rapidly boosts cellular glutathione levels following a single dose in healthy individuals. Efficacy of GGC against *Pseudomonas aeruginosa* derived lipopolysaccharide (LPS), a prominent factor in mediating both bacterial virulence and host responses, in CF remains unassessed.

**Methods:** Primary F508del/F508del mucociliary differentiated bronchial and nasal epithelial cells were created to model LPS-induced oxidative stress and inflammation of CF. The proteomic signature of GGC treated cells was resolved by qLC-MS/MS. Parameters including cell redox state (glutathione, ROS), anti-inflammatory mediators (IL-8, IDO-1) and cellular health (membrane integrity, stress granule formation and cell viability) were assayed.

**Results:** Proteomic analysis identified perturbation of several pathways related to cellular respiration and stress responses upon LPS challenge. Most of these were resolved when cells were treated with GGC. While GGC did not resolve LPS-induced IL-8 and IDO-1 activity, it effectively attenuated LPS-induced ROS and stress granule formation, while significantly increasing intracellular glutathione levels and improving epithelial cell barrier integrity. Moreover, we compared the effect of GGC with thiols NAC and glutathione on cell viability. GGC was the only thiol that increased cell viability; protecting cells against LPS induced cell death. Both therapeutic and prophylactic treatments were successful.

**Conclusion:** Together, these findings indicate that GGC has therapeutic potential for treatment and prevention of oxidative stress related damage to airways in Cystic Fibrosis.

## Introduction

The thiol antioxidant glutathione (GSH) is produced and maintained by all cells of the body at the millimolar range of concentrations. Extracellular GSH concentrations can vary greatly, with plasma levels typically in the low micromolar range. In the epithelial lining fluid (ELF) of the lung, GSH can be 100-times higher than plasma concentrations, with increases occurring in response to stimuli such as pathogens [1, 2]. High GSH concentrations in the ELF, and the ability to raise them, is likely an adaptive response to protect the lung epithelial cells against cytotoxic damage caused by bacterial and viral infections [2]. Transport of GSH out of lung cells is predominantly conducted by the apically expressed cystic fibrosis transmembrane regulator (CFTR) protein [3]. CFTR, which is a member of the ATP-binding cassette (ABC) family of membrane transport proteins, forms a chloride and bicarbonate channel. CFTR thus maintains homeostasis and fluid secretion for many tissues and organs including lung, intestine, and kidney, along with sweat and pancreatic ducts. Unsurprisingly, patients with cystic fibrosis (CF), which is an inherited disease caused by mutations in the CFTR gene, are characterised by a generalised systemic glutathione deficiency and major lung function pathology [4].

CFTR dysfunction deregulates fluid secretion in the lungs, which leads to thick mucus formation and recurrent bacterial infections. Abnormally high levels of reactive oxygen species (ROS), with elevated pro-inflammatory markers are observed in the developing CF foetus even prior to exposure to microorganisms [5]. This innate oxidative state has been attributed to alterations in cellular proteostasis and the mitochondrial electron transport chain caused by deregulated processing and misfolded mutated CFTR. Release of bacterial toxins, mostly derived from the persistent *Pseudomonas aeruginosa* infection, further provokes an exaggerated neutrophilic inflammatory response that produces additional ROS [6]. The uncontrolled production of ROS can be detrimental through the promotion of aberrant cell signalling responses and the oxidative modification of biomolecules leading to lipid peroxidation, protein oxidation and dysfunction, and DNA strand-breaks, that together can promote loss of viability of lung epithelial cells [5]. Under normal physiological conditions, ROS generation is controlled by various adaptive cellular stress responses including the transient formation of cytoplasmic stress granules (SGs), activation of autophagy and alterations to mitochondrial function. ROS generation is balanced by several antioxidant defence systems, which include enzymes such as superoxide dismutase (SOD), catalase, peroxiredoxins (Prx) and glutathione peroxidases (GPXs), which employ the major intracellular reducing agent glutathione (GSH) as a substrate to detoxify ROS [5]. However, CF patients are characterised by GSH deficiency [4] meaning they are exposed to a chronic burden of elevated ROS that cannot be effectively mitigated.

Unsurprisingly, there is a considerable body of research targeting the GSH biosynthesis pathway as a therapeutic strategy to restore GSH homeostasis in CF. Extracellular GSH is unable to be transported into the majority of human cells, as passive uptake of GSH is thermodynamically not feasible due to the high GSH concentration within cells. Therapeutic effects from oral and inhaled formulations of thiols (glutathione and its pro-drug N-acetylcysteine (NAC)) have not demonstrated improvements in clinical outcomes [7, 8]. The immediate precursor to glutathione, γ-glutamylcysteine (GGC), a dietary supplement has shown superior efficacy in increasing intracellular GSH levels in cardiac [9], liver [10] and neural [11] cells *in vitro*. Animal safety trials have demonstrated that GGC is safe at repeated doses at a limit dosage of 1000 mg/kg over a 90-day period [12]. A randomised human pilot study found that single doses of oral administered GGC at 2 g and 4 g significantly increased GSH levels in lymphocytes within 3h, with a return to normal homeostatic levels by 5h [13]. Recently, anti-oxidative and anti-inflammatory properties have been demonstrated for GGC in astrocytes (7, 21) and *in vivo* in a mouse sepsis model (19).

The use of GGC as a potential antioxidant and anti-inflammatory in CF has yet to be explored. The objective of this study was to determine the effectiveness of the GGC against *Pseudomonas aeruginosa* cell-surface lipopolysaccharide (LPS), a prominent factor in mediating both bacterial virulence and host responses.

## Materials and Methods

### Airway epithelial cell culture

Human airway basal epithelial cells were obtained from brushing the nasal inferior turbinate and from bronchoalveolar lavage fluid (BALF) of four CF and two non-CF donors during bronchoscopy. All CF subjects had homozygous DF508-CFTR genotype. The participants’ carers provided written informed consent. This study was approved by the Sydney Children’s Hospital Ethics Review Board (HREC/16/SCHN/120).

#### Conditionally reprogrammed epithelial cell (CREC) culture

Primary airway cultures were established based on a previously published protocol [35]. At confluency, cells were dissociated using a differential trypsin method. Cells were either cryopreserved, or directly utilised for the 2D-conventional or the 3D-differentiated ALI cultures.

#### 3D- Air-Liquid Interface (ALI) differentiated epithelial cell culture

CRCEs were seeded at ~125,000 on Collagen I (Advanced Biomatrix Purecol 5005) coated 6.5 mm 0.4 μM Corning® Transwell® porous polyester membranes (Sigma CLS3470). Cultures were supplemented with PneumaCult Ex-Plus Expansion media (Stemcell 05040) for 4-5 days. Once confluent, the media from the apical side was removed. PneumaCult ALI media (Stemcell 05001) was added on the baso-lateral side. Cultures were monitored for mucociliary differentiation over a period of 21-24 days before experiments were conducted. All experiments were carried out on cells between P1-P2.

#### 2D-conventional epithelial cell culture

CRECs were seeded in a Collagen I coated (Advanced Biomatrix Purecol 5005) 96 well plate and supplemented with Bronchial Epithelial Growth Medium (BEGM) (Lonza™ CC-3170). These cultures were used to assess stress granule formation and cell viability.

### Sample preparation

Matched nasal and bronchial airway CRECs were created from four CF and two non-CF donors. ALI differentiated HBEC cultures from the CF donors (n=4) were cultured and treated under six experimental conditions: mock, LPS^+^, therapeutic, prophylactic, T+P and GGC only with 50μM GGC and 100 μM LPS. The supernatant was saved for IL-8 cytokine and IDO-1 analysis. Cells were harvested in equal aliquots for mass spectrometry and oxidant-antioxidant content analyses. Experiments were performed in triplicate. ALI differentiated HBECs (n=3) and HNEC (n=1) were used for tight junction ZO-1 imaging. With the limitation of primary bronchial epithelial proliferation capacity, assessment of stress granule and cell viability were carried out on the HNECs from the same donors. HNEC from the non-CF donors were used for the cell viability assessment.

### Mass Spectrometry

For the mass spectrometry analysis, total protein was determined by homogenising the cells in RIPA buffer (Life Technologies 89900) containing protease inhibitor cocktail (Sigma 11836153001). Samples were sonicated using the Bioruptor Pico (Diagenode B01060010) for a total of 10 min. Protein concentrations were determined using the 2-D Quant kit (GE Life Sciences 80648356). Samples were reduced (5 mM DTT, 37C, 30 min), alkylated (10 mM IA, RT, 30 min) then incubated with trypsin at a protease:protein ratio of 1:20 (w/w) at 37°C for 18 h, before being subjected to SCX clean-ups (Thermo Fisher, SP341) following manufacturer’s instructions. Eluted peptides from each clean-up were evaporated to dryness in a SpeedVac and reconstituted in 20 μL 0.1% (v/v) formic acid. Proteolytic peptide samples were separated by nanoLC using an Ultimate nanoRSLC UPLC and autosampler system (Dionex, Amsterdam, Netherlands) and analysed on a Tribrid Fusion Lumos mass spectrometer (Thermo Scientific, Bremen, Germany) as described before [36].

### Sequence Database Searches and Protein Quantification

Raw peak lists derived from the above experiments were analysed using MaxQuant (version 1.6.2.10) [37] with the Andromeda algorithm [38]. Search parameters were: ±4.5 ppm tolerance for precursor ions and ±0.5 Da for peptide fragments; carbamidomethyl (C) as a fixed modification; oxidation (M) and N-terminal protein acetylation as variable modifications; and enzyme specificity as trypsin with two missed cleavages possible. Peaks were searched against the human Swiss-Prot database (August 2018 release). Label-free protein quantification was performed using the MaxLFQ algorithm with default parameters [39] and statistical analysis of differential protein abundance were performed in Perseus (version 1.6.5.0) [40]. Functional analysis of differentially abundant proteins was performed with IPA (QIAGEN Inc.)[14]. Mass spectrometry data are available at the ProteomeXchange Consortium via the PRIDE partner repository with the dataset identifier PXD019084 (https://www.ebi.ac.uk/pride/archive/projects/PXD019084). The full list of identified proteins and differentially abundant protein analysis is available upon request.

### Measurement of intracellular glutathione levels

Total intracellular glutathione content (reduced plus oxidised glutathione) was measured using a colorimetric glutathione assay kit (Sigma CS0260), measured by absorbance at 412 nm (Versamax microplate reader Molecular Devices).

### Oxidative Stress Assay

CellROX Green fluorogenic probe was used for measuring oxidative stress levels following manufacturer’s instructions (Life Technologies C10444). The cell-permeable reagents are non- or very weakly - fluorescent while in a reduced state and upon oxidation exhibit strong fluorogenic signal. CellROX® Green Reagent is a DNA dye, and upon oxidation, it binds to DNA; thus, its signal is localized primarily in the nucleus and mitochondria. The fluorescence intensity (485 nm excitation/520 nm emission) was measured with a PerkinElmer EnSight multimode plate reader.

### Enzyme-Linked Immunosorbent Assay (ELISA)

Cytokine IL-8 content in the cell-free culture media, were assessed using an IL-8 ELISA kit (R&D systems DY208) as per manufacturer instruction. Samples were diluted 1/10 and all values fell within the standard curve. The ELISA plate was read using the Versamax microplate reader (Molecular Devices).

### High performance liquid chromatography (HPLC)

IDO-1 dioxygenase activity was determined by measuring the extent of conversion of L-tryptophan (L-Trp) into kynurenine (Kyn) in the culture medium using an Agilent-1260 HPLC system. For this, the cell-free culture medium was treated with 20% trichloroacetic acid in a 3:1 (v/v) ratio to precipitate proteins and centrifuged (10 minutes, 18000 g, 4°C). Protein free supernatants were then injected into a Hypersil 3-μm ODS-C18 column (Phenomenex, USA) and eluted at 0.5 mL/min with a mobile phase consisting of 9% (v/v) acetonitrile and 100 mM chloroacetic acid (pH 2.4). L-Trp and Kyn were detected at 280 and 364 nm, respectively and their concentrations determined by peak-area comparison of L-Trp and Kyn peaks in experimental samples with authentic L-Trp and Kyn standards of known concentration. IDO-1 activity was presented as the Kyn:L-Trp ratio.

### Immunofluorescence and imaging

Cells were fixed in 3.7% formaldehyde (Thermo Scientific 28906) followed by neutralisation (100 mM glycine for 10 min at room temperature) and permeabilization (0.5% Triton-X for 10 min. The cells were blocked with 10% goat serum (Sigma G9023) for 90 min at room temperature prior to overnight incubation with primary antibodies (**Table S1**). Incubation with fluorescently-conjugated secondary antibodies was carried out for 1 h. Cells were mounted with Vectashield hardset antifade medium with DAPI (Vector Laboratories H-1500). For tight junction immunostaining, methanol-acetone fixation for 15 minutes at −20’C was performed.

Imaging to validate the cell models was performed using Leica TCS SP8 DLS confocal microscope using a 20X/0.75 HC PL APO CS2 air objective lens. Stress granule images were acquired via a Zeiss LSM 780 inverted laser scanning confocal microscope using a 20X/0.8 Plan-Apochromat air objective lens. Quantification of the percentage of cells with stress granules was measured by manually counting the number of cells with ≥3 visible foci per cell. To characterize ZO-1 expression, z-stacks of multiple fields of view (47-55 per treatment) were acquired using Leica TCS SP8 DLS confocal microscope, 63x/1.4 HC PL APO CS2 oil immersion objective. Images were analysed using custom-built script in Matlab (MathWorks, Natick, MA).

### Cell viability/proliferation assay

Thiazolyl blue tetrazolium bromide (MTT) (Sigma M5655) was added to a final concentration of 0.5 mg/mL. Cells were incubated for 2 h under 5% CO_2_ at 37°C prior to the addition of MTT and 200 μl DMSO to solubilise the cells. The absorbance of each well was measured at 570 nm using a Versamax microplate reader (Molecular Devices).

### Antioxidant compounds

GGC sodium salt (Biospecialties International, Mayfield, NSW, Australia), NAC (Sigma A7250) and GSH (Sigma G4251) were prepared fresh prior to each experiment in cold PBS (Sigma 806552).

### Statistical analysis

The data are presented as a mean with errors representing standard deviation (SD). Single group statistical analysis was performed using one-way ANOVA, while the two group analysis used two-way ANOVA. All statistical calculations were performed using GraphPad Prism, with the exception of the aforementioned analyses of quantitative proteomics data. P values <0.05 were considered statistically significant with higher degrees of significance described in figure legends.

## Results

We established LPS-challenged 3D-differentiated human bronchial and nasal epithelial cells models and confirmed that they exhibit a dose-dependent increase in ROS production, while producing significant amounts of the pro-inflammatory cytokine IL-8 (**Figure S1**). Cells were treated under six experimental conditions: (1) PBS control (Mock), (2) LPS-challenged (LPS^+^), (3) GGC added 24 h post-LPS challenge (therapeutic, T), (4) GGC added 24 h prior to LPS challenge (prophylactic, P), (5) GGC added 24 h prior and 24 h post-LPS challenge (T+P); and (6) GGC alone (**Figure 1**).

**Figure 1.**
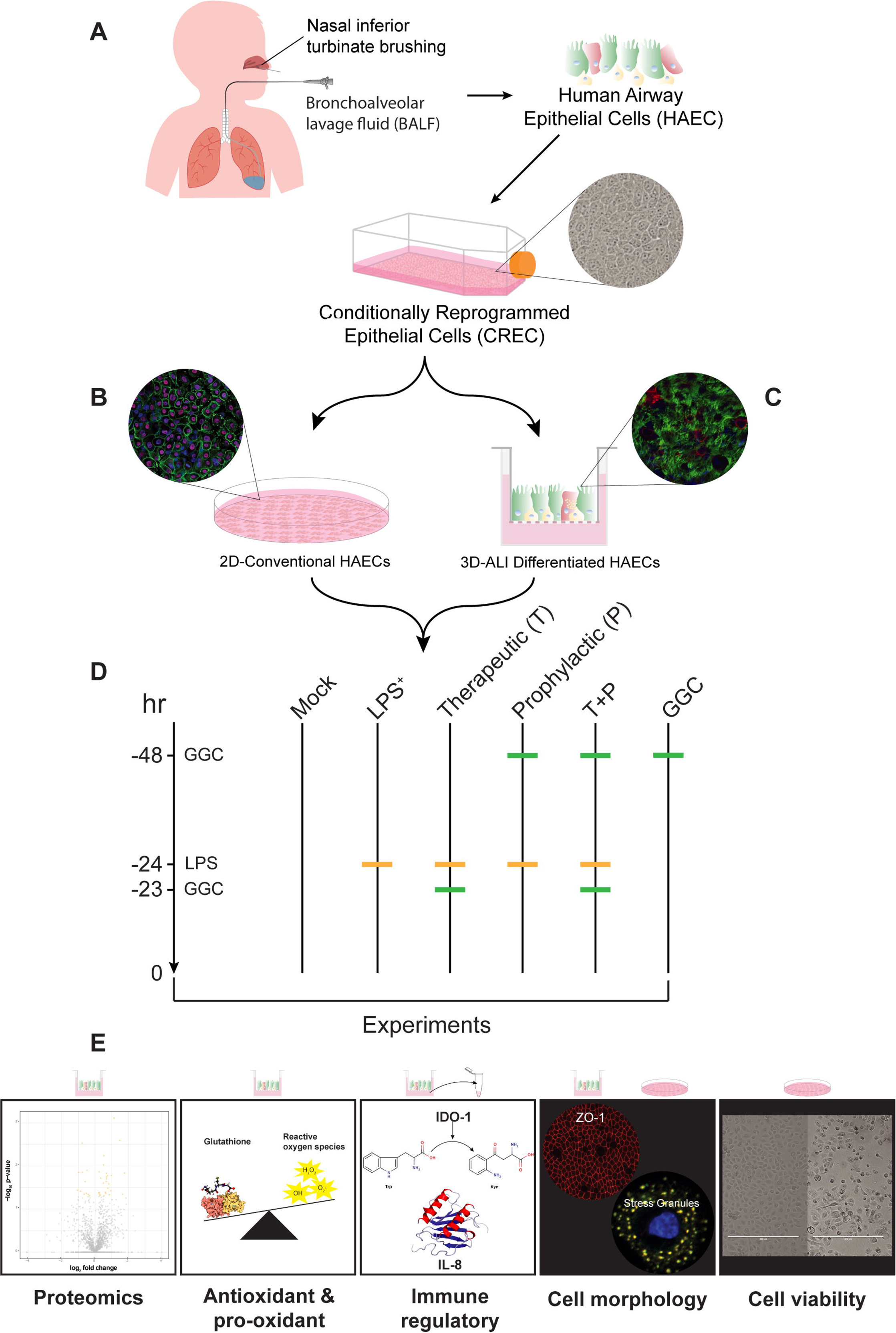
Schematic of experimental protocol. **(A)** Airway basal epithelial cells from four CF donors were used to create conditionally reprogrammed airway epithelial cells (CREC). CRECs were established in **(B)** 2D-conventional epithelial monolayers or in **(C)** 3D terminally differentiated pseudostratified epithelium at Air Liquid Interface (ALI). **(D)** Treatments used to explore the potential efficacy of GGC against *Pseudomonas aeruginosa* derived lipopolysaccharide (LPS) included: (1) PBS control (Mock), (2) LPS-challenged (LPS^+^), (3) GGC added 24 h post-LPS challenge (therapeutic, T), (4) GGC added 24 h prior to LPS challenge (prophylactic, P), (5) GGC added 24 h prior and 24 h post LPS challenge (T+P); and (6) GGC alone **(E)** Proteomics, antioxidant and pro-oxidant, inflammatory response, cell morphology and cell viability analyses were carried out 24 h post infection for each treatment condition in primary differentiated human airway epithelial cells (HAEC) of either bronchial (HBEC) or nasal (HNEC) origin.

### LPS challenge or GGC treatment alters protein expression in differentiated primary human bronchial epithelial cells

We first investigated the extent of alteration of the human bronchial epithelial cells (HBEC) proteome in response to LPS challenge and GGC treatment. Between 1714 and 1944 proteins were identified with quantitative LC-MS/MS in each of the six treatment conditions (**Table S2)**. To identify significantly differentially abundant proteins, pairwise comparisons of conditions were performed. The three GGC treatments (therapeutic, prophylactic, T+P) were compared against the LPS-challenged proteome. LPS-challenged and GGC only proteomes were compared to that of the mock control (**Figure 2, Table S2**). In the LPS-challenged comparison, the tight junction protein ZO-1 and immune regulatory or inflammatory proteins (IDO-1, S100A2, CXCL6 and MUC5B) were amongst the most differentially abundant (**Figure 2A**). In both the therapeutic (**Figure 2B**) and prophylactic (**Figure 2C**) comparisons, NAD(P)H-dependant oxidoreductase proteins (BDH2, NDUS3, NDUA8 and AL4A1), the pro-survival protein TCTP, antioxidants peroxiredoxin 3 (PRDX3) and ROS scavenger SERPINB5 were significantly more abundant. Glutathione peroxidase 4 (GPx4), which uses glutathione to protect cells against lipid peroxidation, was significantly more abundant in the therapeutic comparison (**Figure 2B**) condition. In the T+P comparison, three oxidative stress responsive proteins (CIRBP, PARP1 and APEX1) were differentially abundant (**Figure 2D**). The antioxidant enzyme, glutaredoxin (GLRX), which employs glutathione as a cofactor, was the most significantly altered protein in the comparison of cells treated with GGC alone (**Figure 2E**).

**Figure 2.**
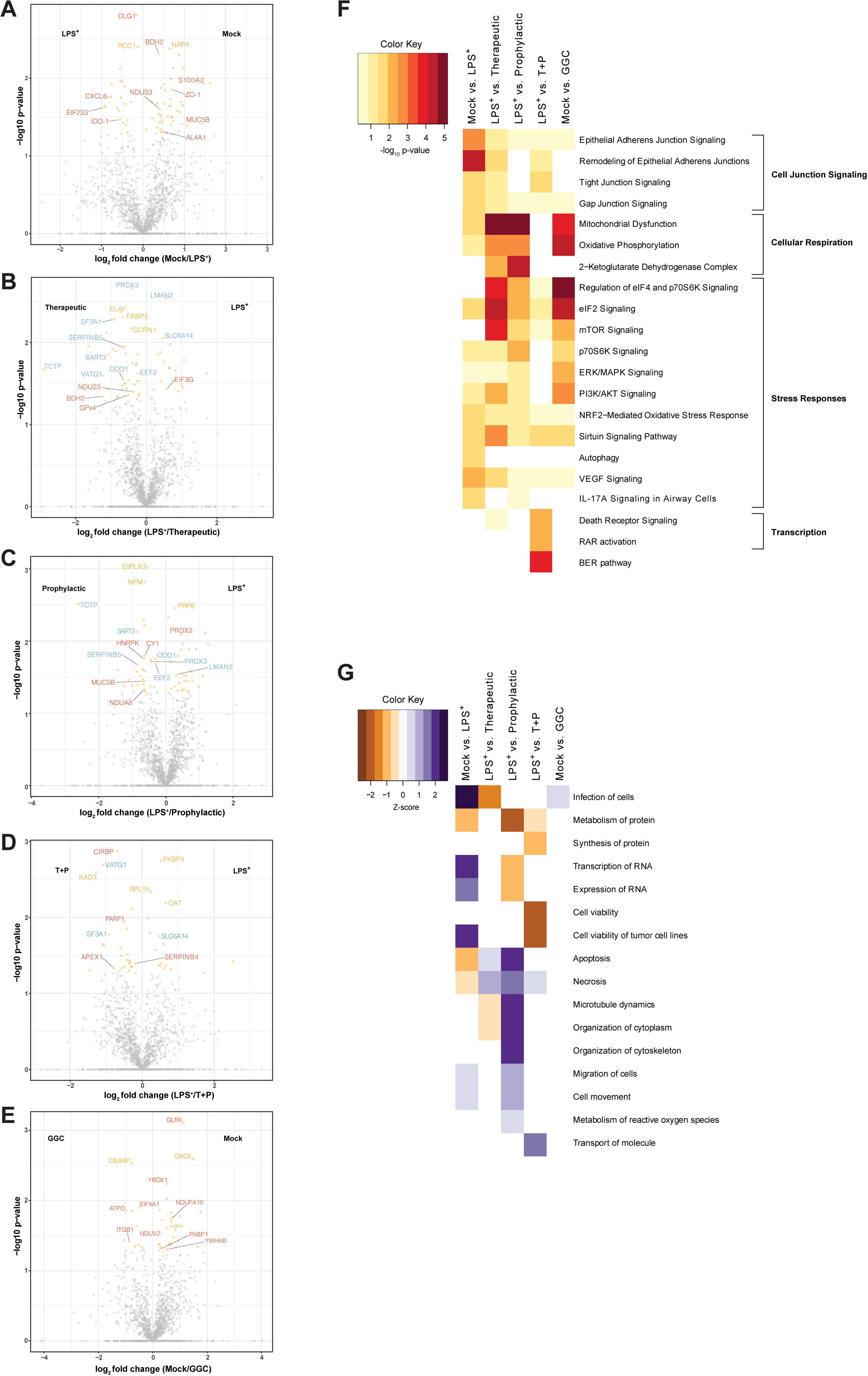
Differential protein abundance in differentiated human bronchial epithelial cells after LPS challenge and GGC treatment. Proteomes of cells from mock, LPS^+^, therapeutic (T), prophylactic (P), T+P and GGC only conditions were assessed. Volcano plots of differentially abundant proteins between pairwise comparisons of: **(A)** mock and LPS^+^, **(B)** LPS^+^ and therapeutic treatment, **(C)** LPS^+^ and prophylactic treatment, **(D)** LPS^+^ and T+P treatment, and **(E)** mock and GGC only conditions. The log_10_ transformed p-values (y-axis) were plotted against the average log_2_ fold-change (x-axis). Proteins represented by orange dots were statistically significant (p < 0.05) and grey dots were not statistically significant. Proteins of biological interest (red) and differentially abundant in multiple comparisons (blue) are labelled. **(F)** A heatmap of the enriched canonical pathways among differentially abundant proteins in the five comparisons (A-E) discovered by Ingenuity Pathway Analysis (IPA). Colour indicates −log_10_ p-value of pathways with high statistical significance (red) and low statistical significance (yellow). **(G)** Heatmap of enriched disease biological functions among differentially abundant proteins in the five comparisons (A-E) discovered by IPA. Colour indicates the activation Z-score and predicts whether a disease or biological function is increased (positive Z-score; blue) or decreased (negative Z-score; brown) based on the experimental dataset. Data are from four CF derived differentiated HBECs with three technical replicates for each different experimental condition.

### Biological pathways involved in cellular respiration, stress response and cell junction signalling are altered following the GGC-treatment of LPS-challenged cells

Ingenuity pathway analysis (IPA) [14] of canonical pathways, networks and biological functions of differentially abundant proteins in each comparison pair are presented in **Figure 2F** and **Table S4**. Altered canonical pathways in the LPS-challenged comparison include oxidative and inflammatory responses such as NRF2-mediated oxidative stress signalling and the IL-17A cytokine signalling pathway. In both therapeutic and prophylactic treatment comparisons, enriched canonical pathways were related to cellular respiration, protein synthesis and stress adaptation responses. These same pathways were enriched in the cells treated with GGC alone. Enriched pathways in the T+P condition were involved in the prevention of apoptosis, activation of transcription and stress response (**Figure 2F**).

Disease and biological function analysis predicted decreased *cell viability* in LPS-challenged cells. In contrast, *cell viability* was predicted to increase in the T+P treatment. *Apoptosis* and *necrosis* were predicted to decrease in the prophylactic treatment (**Figure 2G**).

### GGC increased the glutathione (GSH) level and decreased ROS production in LPS-challenged differentiated primary human bronchial epithelial cells

Proteomic analysis revealed increased abundance of several antioxidant enzymes in HBECs with GGC treatment (**Figure 2 and Table S3**). We therefore assessed intracellular GSH levels in the cells from the six different treatment conditions (**Figure 3A**). LPS treatment significantly lowered intracellular GSH levels by 50% compared to mock control cells. LPS-induced GSH depletion was ameliorated by all GGC treatment conditions. Intracellular GSH was restored to near unchallenged basal GSH levels. The LPS-induced reduction in GSH levels coincided with a marked increase in cellular ROS production (**Figure 3B**). This LPS-induced increase in ROS was significantly decreased with therapeutic, prophylactic and T+P GGC treatments. (**Figure 3B**).

**Figure 3.**
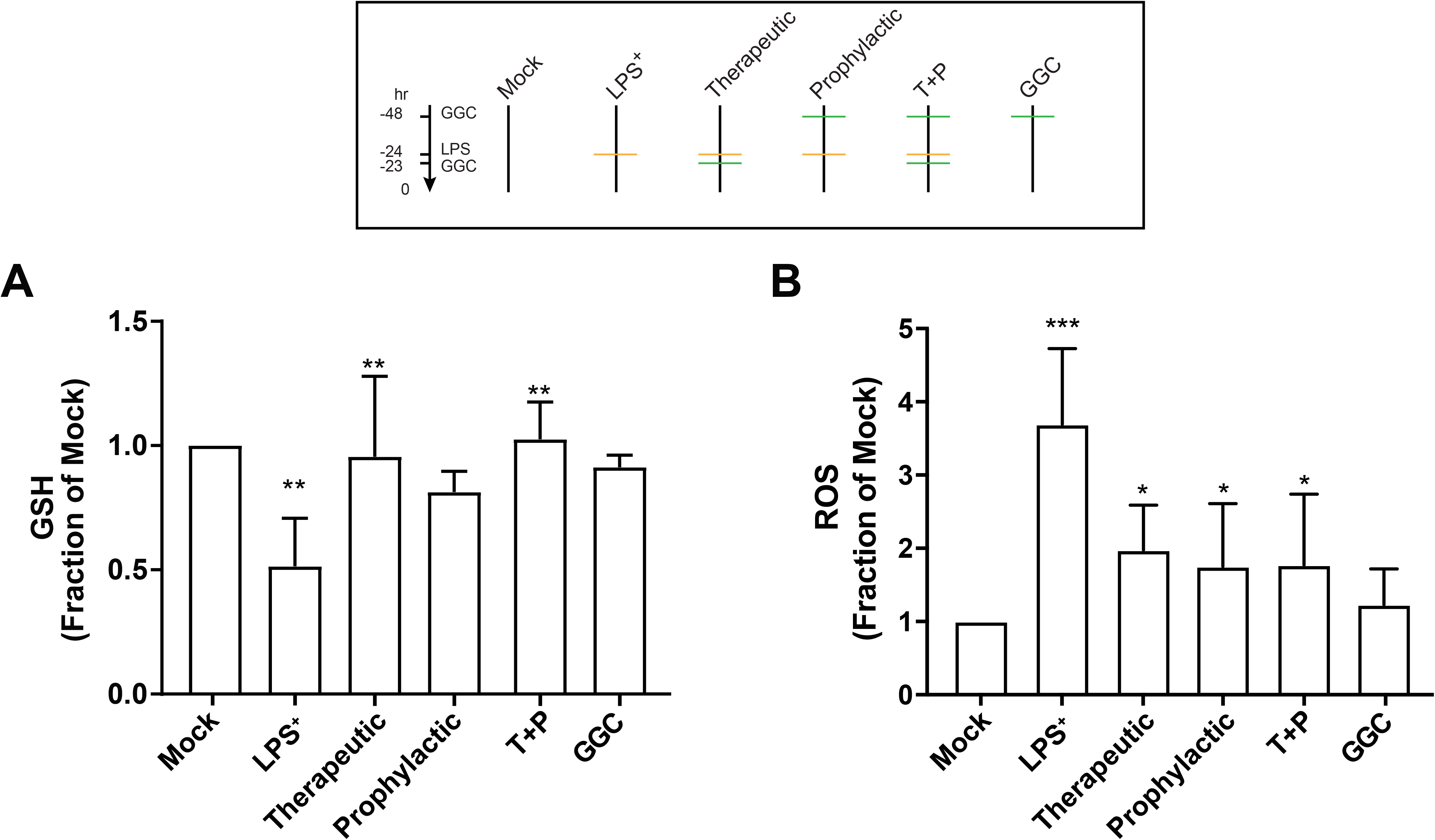
GGC ameliorates LPS-induced reduction in endogenous glutathione levels and increase in cellular Reactive Oxygen Species (ROS) levels. Cells from mock, LPS^+^, therapeutic (T), prophylactic (P), T+P and GGC only conditions were assayed for **(A)** total intracellular glutathione (GSH) and **(B)** ROS levels. Statistical analysis was performed using two-way ANOVA against LPS^+^ for therapeutic, prophylactic and T+P conditions, and mock for LPS^+^ and GGC only (* = p<0.05, ** = p<0.01, *** = p<0.001). Error bars represent SD. Data are from four CF derived differentiated HBECs with three technical replicates for each different experimental condition.

### GGC did not alter the inflammatory and immune regulatory responses in LPS-challenged differentiated primary human bronchial epithelial cells

IDO-1, a L-tryptophan catabolizing immune regulatory enzyme that is induced during inflammation [15], was significantly up-regulated in the proteome of the LPS-challenged cells (**Figure 2A**). Additionally, the majority of differentially abundant proteins were involved in *antimicrobial response*, and *inflammatory response* networks (**Table S4**). In comparison, the three GGC treated conditions did not reveal enrichment in these networks (**Table S4**). IDO-1 and pro-inflammatory cytokine IL-8 were assessed to determine whether GGC can alleviate LPS-induced inflammation. LPS challenge resulted in a four-fold increase in IL-8 levels (**Figure 4A**). A 24-fold increase in cellular IDO-1 enzyme activity (indexed as the kynurenine/tryptophan ratio) was also observed in LPS-treated cells. However, treatment with GGC did not alter the LPS-induced increases in either IL-8 levels (**Figure 4A)**or IDO-1 activity **(Figure 4 B**).

**Figure 4.**
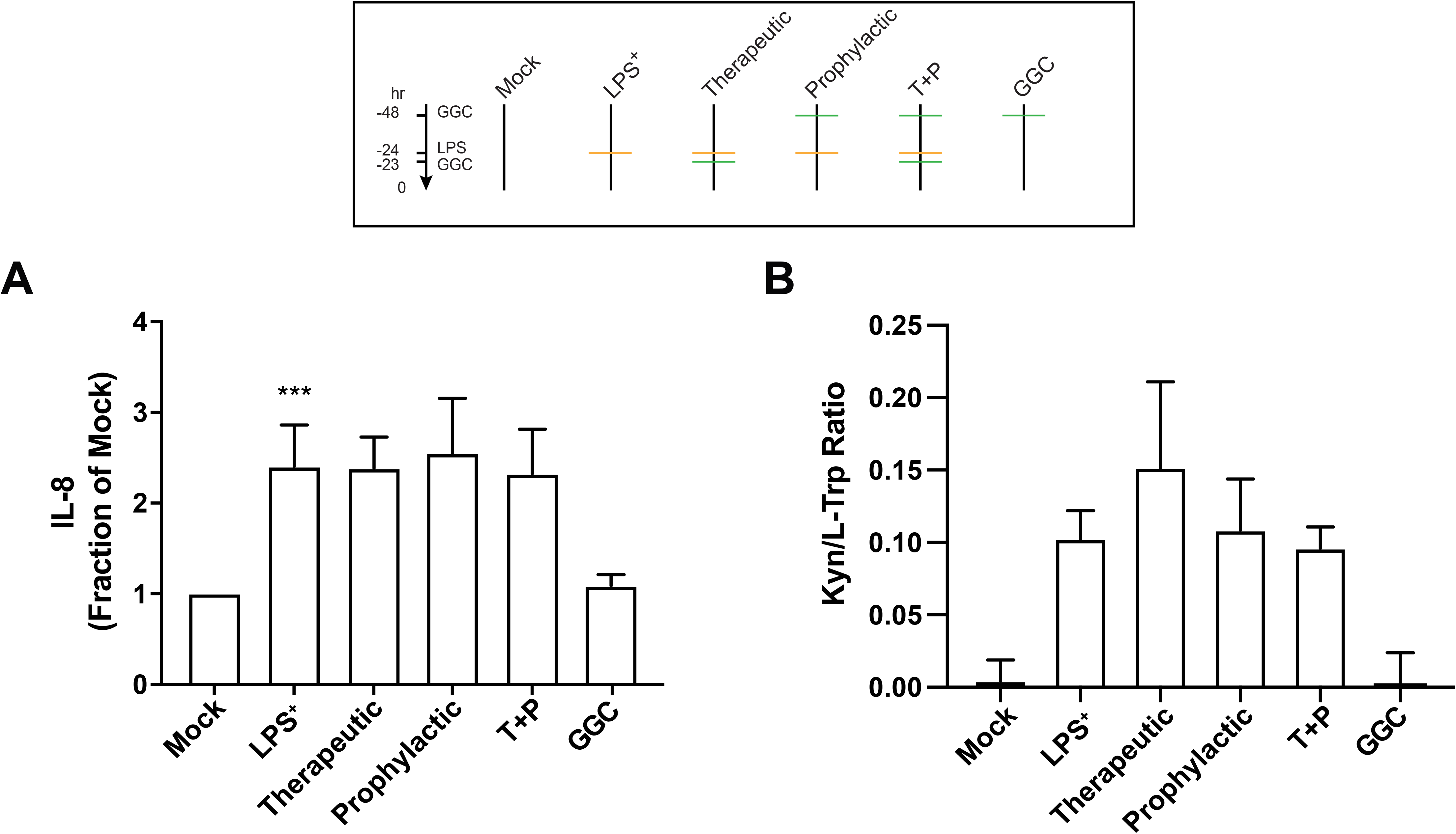
GGC does not abrogate LPS-induced inflammatory and immune regulatory responses. Culture media collected from the proteomics analyses from mock, LPS^+^, therapeutic (T), prophylactic (P), T+P and GGC only conditions were assayed for (**A**) IL-8 cytokine and (**B**) IDO-1 activity measured as the kynurenine to tryptophan ratio. Statistical analysis was performed using two-way ANOVA against LPS^+^ for T, P and T+P conditions, and mock for LPS^+^ and GGC only (*** = p<0.001). Error bars represent SD.

### GGC alleviated LPS-induced deterioration of epithelial tight junction protein zona occludin-1 (ZO-1) in human bronchial and nasal epithelial cells

Proteomic analysis indicated a 44% decrease in epithelial tight junction protein ZO-1 expression in the proteome of the LPS-challenged cells (**Figure 2A)**. Furthermore, tight junction and gap junction signalling pathways were enriched in LPS-challenged condition compared to mock (**Figure 2F, 5A**). We confirmed this observation by immunofluorescence staining for ZO-1 expression in differentiated bronchial and nasal epithelial cells (**Fig 5B**). LPS-challenge significantly reduced the protein expression of ZO-1 in the HAECs compared to the unchallenged control cells (**Figure 5D**). All three GGC treatments ameliorated the LPS-induced reduction in ZO-1 expression. ZO-1 levels were increased above the basal level in therapeutic and T+P. Treatment with GGC in the absence of LPS did not alter ZO-1 expression (**Figure 5C-D**).

**Figure 5.**
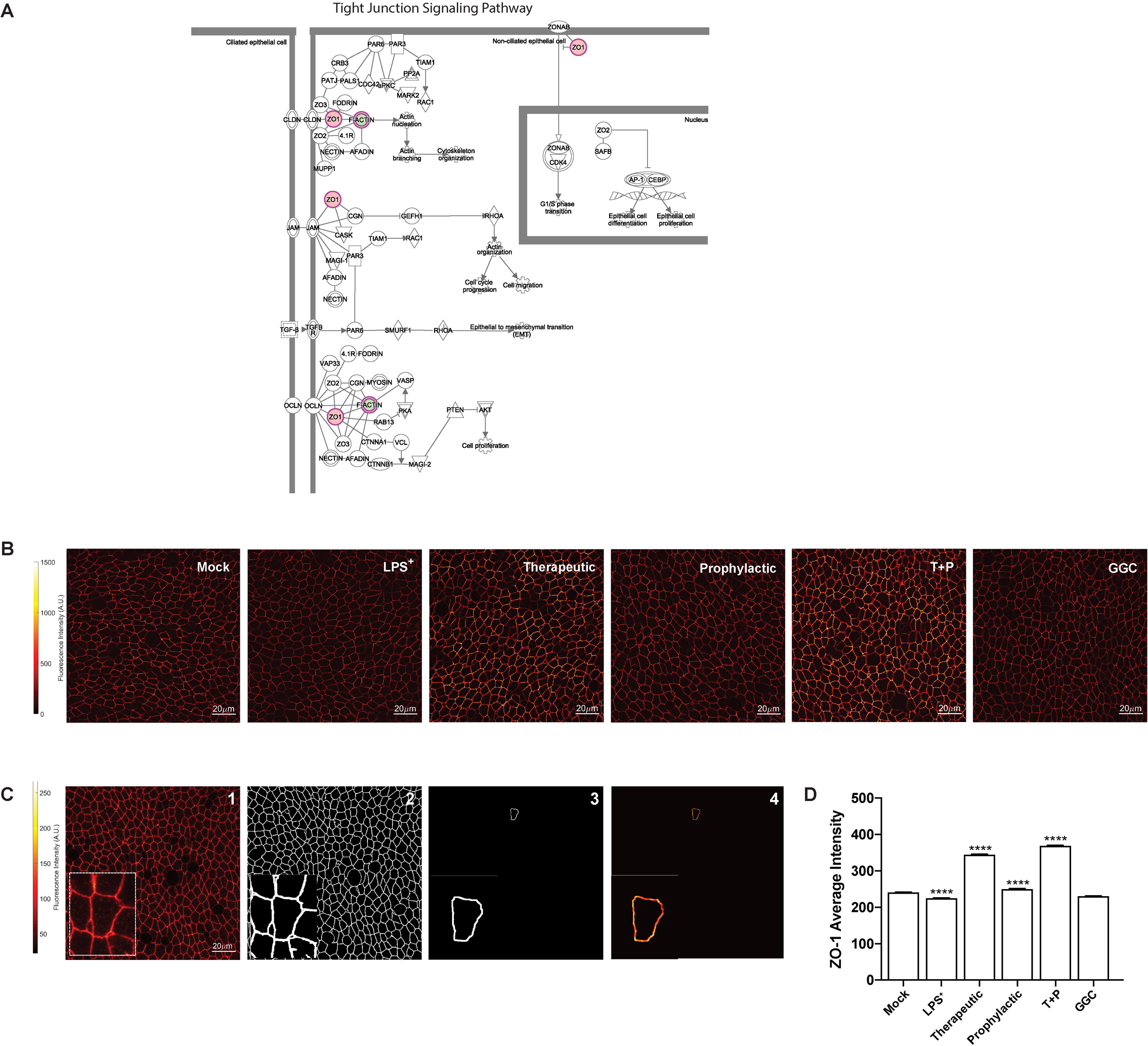
Reduction of tight junction protein ZO-1 by LPS challenge was ameliorated with GGC. Mock, LPS^+^, therapeutic (T), prophylactic (P), T+P and GGC only conditions were assessed for tight junction protein ZO-1 expression. **(A)** Tight junction signalling pathway. Red highlighted proteins were more abundant in LPS^+^ relative to the mock, while green was less abundant in LPS^+^ relative to the mock. The pathway was adapted from IPA (QIAGEN). **(B)** Representative image of ZO-1 immunofluorescence in the six experimental conditions. **(C)** Summary workflow of ZO-1 quantification. The mid-plane of acquired z-stack images (plane with highest intensity) were determined (panel 1) and used to generate the binary mask of tight junctions (panel 2). The junction of individual cells was extracted (panel 3) and the mid-plane intensity of pixels spanning the entire cellular junction was used to calculate the average intensity and standard deviation of intensity within each cell (panel 4). **(D)** Mean ZO-1 intensity as calculated using the above workflow for four differentiated HAECs (HBEC =3; HNEC =1). Statistical analysis was performed using one-way ANOVA against LPS^+^ for T, P and T+P conditions, and mock for LPS^+^ and GGC only (**** = p<0.0001). Error bars represent SD. Scale bars represent 20 μm.

### GGC prevents LPS-induced stress granule formation in human nasal epithelial cells

Protein synthesis pathways involved in stress response adaptation were enriched in the HBECs treated with GGC (**Figure 2F, 6A**). Differentially abundant proteins in these pathways included G3BP, EIF2S3, EIF3G, and EIF4A1 (**Figure 2**). These proteins are well-characterised markers of cytosolic stress granules (SG). These granules form when translation initiation is stalled [16]. With the limitation of primary bronchial epithelial proliferation capacity, assessment of stress granule formation was carried out in from the same donors’ nasal epithelial cells (HNEC). To confirm whether SG formation is modulated by GGC, we assessed the co-localisation of two SG protein markers, G3BP and EIF4A, in cells (**Figure 6B, S4**). G3BP and EIF4A were seen in small fraction of cells in all conditions, but they only co-localised in the LPS-challenged cells at low frequency (22% ± 8% cells). Therapeutic treatment of GGC completely ameliorated SG formation. Prophylactic and T+P treatment also prevented LPS-induced SG formation.

**Figure 6.**
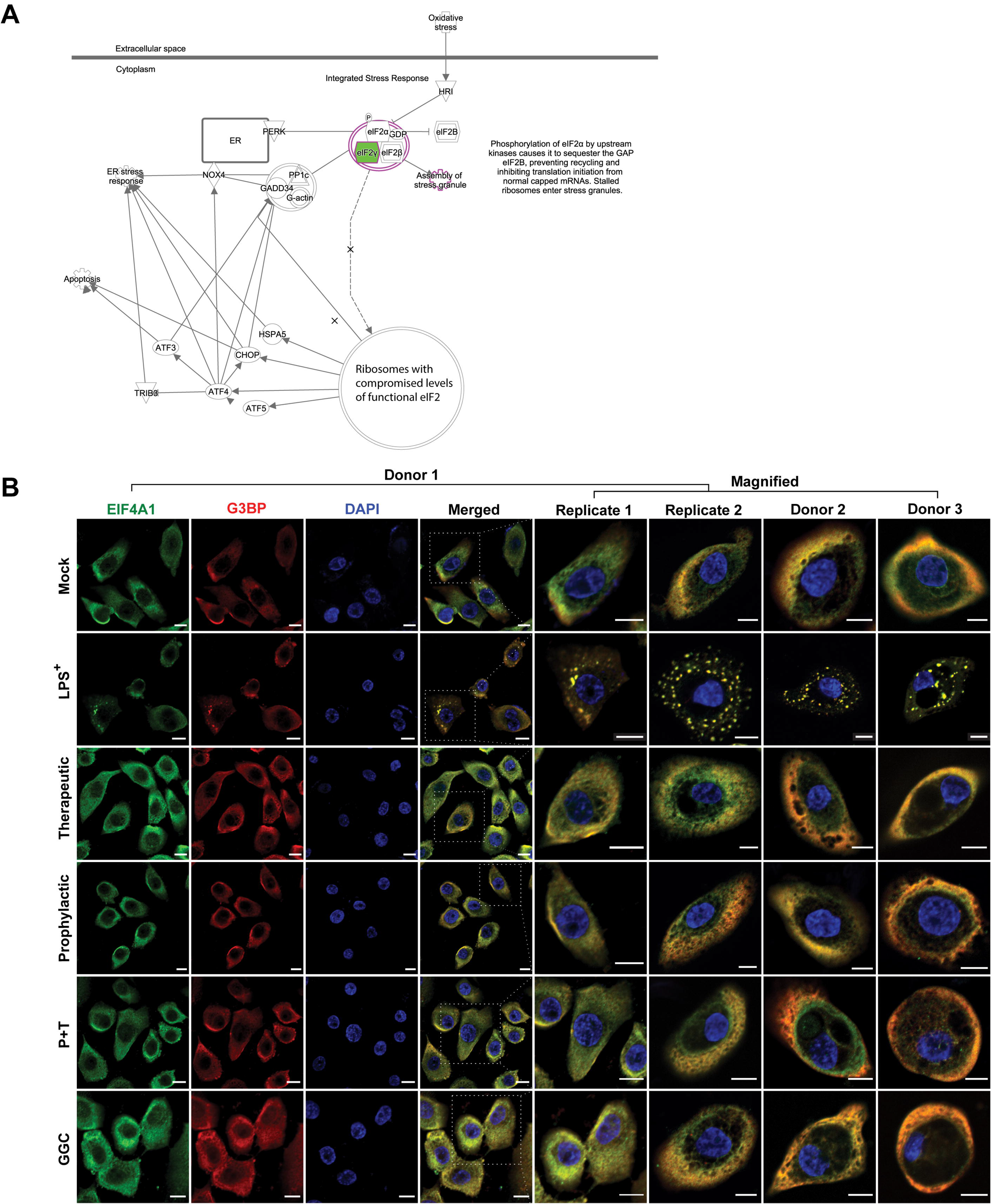
GGC inhibited LPS-induced stress granule formation. Mock, LPS^+^, therapeutic (T), prophylactic (P), T+P and GGC only conditions were assessed for tight junction protein ZO-1 expression. (**A**) The EIF2 signalling pathway, which is involved in stress response adaptation including the assembly of stress granules. Green highlighted proteins were less abundant in LPS^+^ relative to the mock (Figure 3A). The pathway is adapted from IPA (QIAGEN). **(B)** Representative immunofluorescence co-staining for stress granule markers EIF4A and G3BP images from three CF donors 2D HNECs in the six experimental conditions. HNEC cells from Donor 1 were cultured and assessed on two separate occasions (Replica 1 and 2). Stress granules were identified in cells which both EIF4A (green) and G3BP (red) signals overlapped. Nuclei are stained blue. Scale bars represent 10 μm.

### Treatment with GGC increases cell viability in LPS-challenged human nasal epithelial cells

Our proteomic analysis revealed modulation of mitochondrial proteins including NADPH-dependant oxidoreductases (**Figure 2A**), which are enzymes that play central role in cellular metabolism and catalyse various redox reactions [17]. Assessment of NADPH-dependent oxidoreductase activity is commonly carried out with artificial dyes (Formazan) to determine the metabolic activity of cells, and as an indicator of cell viability. LPS challenge significantly decreased NADPH oxidoreductase activity, and consequently cell viability (**Figure 7A**). We assessed primary HNECs and immortalised HBECs treated with GGC, NAC and GSH. Thiol only, therapeutic and T_P treatment conditions were tested (**Figure 7B-D and Figure S2**). Treatment with GGC between 2 to 50 μM significantly increased cell viability in all treatment conditions at the same concentration, NAC and GSH treatment were ineffective in increasing cell viability.

**Figure 7.**
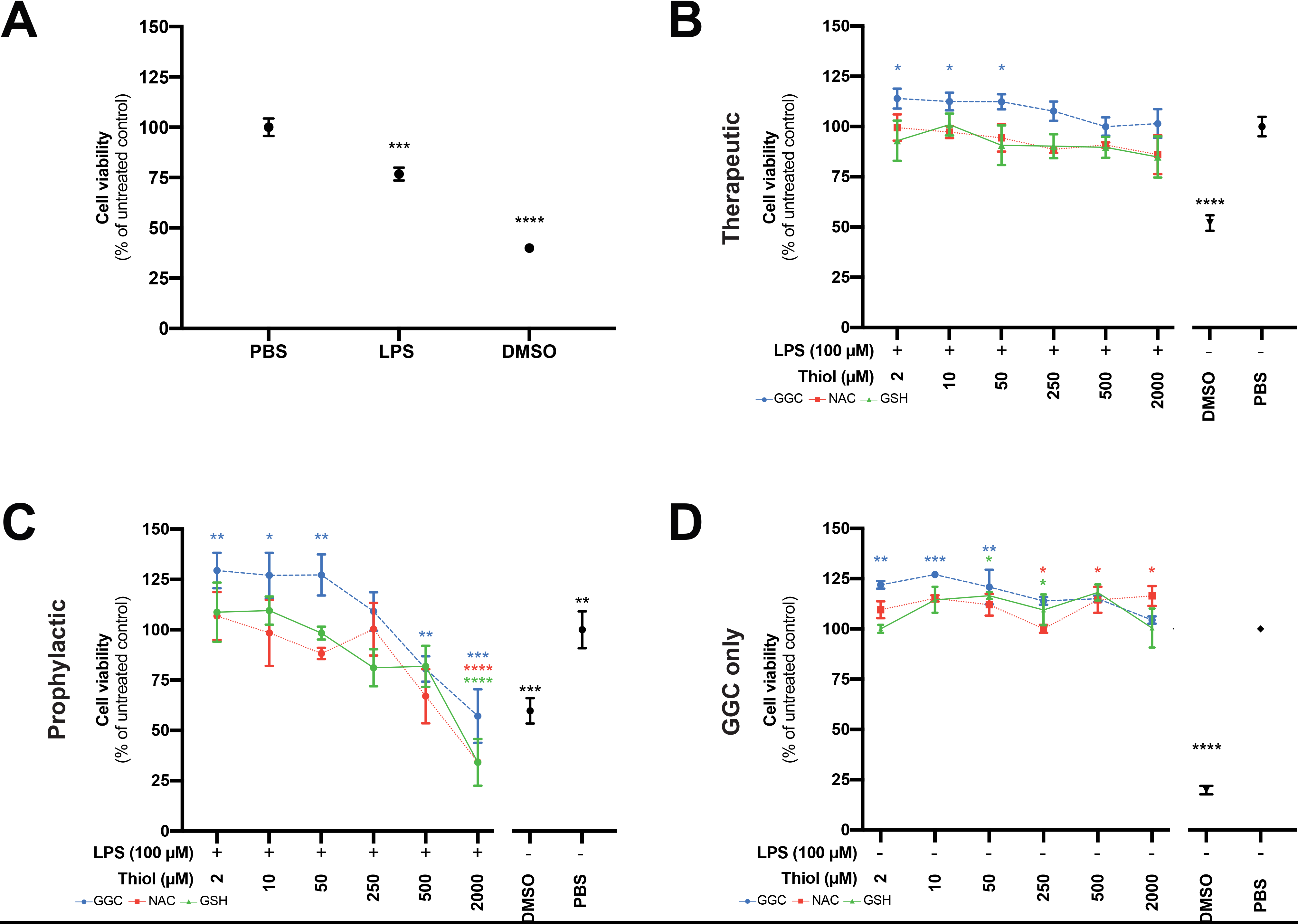
GGC increases cell viability of unchallenged and LPS-challenged human nasal epithelial cells. Cell viability was assessed in (**A)** LPS-challenged untreated (LPS^+^), (**B**) unchallenged treated, **(C)** post-LPS^+^ treated (therapeutic) and (**D**) pre & post LPS^+^ treated (T+P) conditions in CF HNEC using the MTT assay. GGC, NAC and GSH treatment in a range of concentration (2 to 2000μl) were tested. DMSO used as positive cell death control. Statistical analysis was performed using one-way ANOVA against the PBS control. (* = p<0.05, ** = p<0.01, *** = p<0.001, **** = p<0.0001). Error bars represent SD. Data are from three CF HNECs with three technical replicates for each different experimental condition.

## Discussion

Using primary CF airway epithelial differentiated cell models, this study has established that the administration of exogenous γ-glutamylcysteine (GGC) can increase intracellular glutathione (GSH) levels and protect cells from LPS-induced cellular damage. Proteomic analysis identified perturbation of several pathways related to cellular respiration, transcription, stress responses and cell-cell junction signalling upon LPS challenge which were altered when cells were treated with GGC. We confirmed GGC improves tight junction activity and attenuates LPS-induced stress granule formation. Unlike, NAC and GSH, GGC in both therapeutic and prophylactic treatment increased metabolic activity leading to increased cell viability. Our data suggest that GGC therapy could mitigate the lung tissue damage suffered by CF patients from repeated *P. aeruginosa* infections.

Cystic fibrosis is characterised by persistent inflammation, oxidative stress of the airways and deficiency in the major intracellular antioxidant GSH [18]. Despite multiple clinical trials with current glutathione-based therapies, improved clinical outcomes in CF have not been achieved [6–7]. As a precursor to GSH, GGC has been consistently shown to increase intracellular levels of GSH in both *in vivo* and *in vitro* models [9–11, 13, 19–21], and to reduce inflammation and oxidative stress [22–24]. Recently, it was shown that GGC exhibits anti-inflammatory effects in an *in vivo* and *in vitro* mouse sepsis model [19]. However, in our HBEC CF model, administration of GGC had no impact on reducing the LPS-challenged up-regulation of two inflammatory markers, IL-8 and IDO-1. While inflammation and oxidative stress are tightly linked processes that can induce one or the other, they can also occur independently [25–27].

Indeed, GGC administration significantly increased endogenous GSH levels in our LPS-challenged HBEC CF model. Both prophylactic and therapeutic treatment with GGC were effective, with a combination of both being the most effective measure. This concurred with a previous study that showed GSH supplementation ameliorates GSH depletion caused by LPS in an acute lung injury mouse model [28]. Furthermore, GGC proved to be an effective prophylactic and therapeutic treatment for attenuating LPS-induced ROS production.

Endogenous GGC itself is an antioxidant by acting as an alternative co-factor for glutathione peroxidase (GPx), regardless of changes to the intracellular glutathione concentration [22]. We found elevated GPx4 expression in LPS-challenged cells that had been treated with GGC. GPx4 is an essential antioxidant protein that protects against oxidative stress by degrading the ROS hydrogen peroxide (H_2_O_2_) and lipid hydroperoxides [29, 30]. Peroxidation of membrane lipids can disturb the assembly of cell membranes, which inevitably will impact membrane permeability. Intestinal malabsorption of fat soluble vitamins, a characteristic of CF, causes deficiency of the major lipid membrane antioxidant, vitamin E, leading to excessive lipid peroxidation [5]. Further impairment of the membrane integrity is evident by dislocation of epithelial tight junction protein ZO-1 from the membrane to the cytosol and to the nucleus in CF epithelial cells [31]. ZO-1 expression is further reduced with LPS challenge [32]. Downregulation of ZO-1 in LPS-challenged CF cells was evident in both our proteomics and immunofluorescence analyses. Glutathione is essential for maintenance of membrane integrity [29]. GGC treatment post-LPS challenge showed a significant reestablishment of ZO-1 expression at the cell boundaries. Pre-treatment with GGC protected against LPS-induced ZO-1 reduction, though not to the same magnitude as that observed for the therapeutic or pre-and post-GGC treatment. This suggested that treatment with GGC in either prophylactic or therapeutic capacity may be effective for reversing LPS-induced and persistent CF-related membrane integrity impairment, protecting against viral or bacterial entry and spread to other cells.

Inhibition of protein synthesis is a well characterised consequence of exposure of cells to oxidative stress [16]. We observed alterations in protein synthesis pathways, including the EIF2 signalling pathway, in the proteomes of CF HBECs. Inhibition of translation initiation leads to the accumulation of stalled translation pre-initiation complexes (PICs) that condense to form non–membrane-enclosed foci known as stress granules (SGs). SG formation can be triggered by protein misfolding. No SGs were observed in the mock or control cells, which suggested that misfolded mutant CFTR does not cause chronic SG formation. LPS challenge is known to stimulate SG formation [33]. In our model, exposure to LPS caused SG formation in approximately 20% of cells. Elevated levels of GSH has been shown to inhibit arsenite-induced SG formation in West Nile virus-infected cells [34]. Our data demonstrated that GGC effectively inhibits LPS-induced SG formation in CF cells in the therapeutic, prophylactic and combination treatment regimens. The inhibition of SG formation supports that elevation of GSH levels by GGC treatment protects against LPS-induced inhibition of translation initiation, thereby preserving cellular protein synthesis.

Protein synthesis, one of the most energy consuming processes in the cell is a key process for cell viability. We observed protection against LPS-induced drop in metabolic activity, an overall improvement in cell viability, in all GGC treatments in bronchial and nasal cells. Even in absence of LPS, treatment with GGC improved metabolic activity and cell viability. In contrast, NAC and GSH supplementation did not demonstrate significant improvements at the same dose, nor was a consistent and reproducible protective effect observed at higher doses. This observation may explain the lack of success for GSH and NAC clinical trials in improving pulmonary function [7, 8]. We note that treatment at exceptionally high concentration of 2000 μM with all three thiols compromised cell viability in the therapeutic and prophylactic combined condition. With regards to GGC, the 2000 μM equate to 17.5 g of an *in vivo* dose. Animal safety trials have demonstrated that GGC is safe at repeated doses at a limit dosage of 1000 mg/kg over a 90-day period [12]. No adverse effect was observed in a human clinical trial of healthy, non-fasting subjects [13]. Single dose of oral administered GGC at 2 g and 4 g is bioavailable and can increase intracellular GSH levels above homeostasis in lymphocytes.

Our data showed that GGC treatment of CF human airway cells increases the overall cellular redox status of the cell in favour of a less oxidative state, which may alleviate LPS-induced cell stress and ROS production. This may occur through multiple interplaying mechanisms: 1) increasing endogenous GSH levels which directly scavenge ROS; 2) acting as a reducing co-factor for antioxidant proteins (e.g. GPx4); and 3) up-regulating oxidative stress response proteins (e.g. peroxiredoxins). We provide promising data in support of a beneficial effect of GGC for CF antioxidant therapy. GGC has self-affirmed Generally Recognised as Safe (GRAS) status, which should facilitate and simplify its regulatory pathway to the clinic. Products containing GGC for oral consumption are now on sale to the general public in the USA. An oral route of delivery allows for rapid first pass metabolism of GGC via the gut and liver and a subsequent increase in circulating GGC [13]. Defects in gut and liver function in CF patients, however, could hamper oral GGC efficacy. Controlled clinical studies are now needed to investigate GGC’s safety in CF patients and its potential as a useful preventative and therapeutic treatment for CF airway redox imbalance.

## Abbreviations

ABC: ATP-binding cassette
ALI: Air-liquid interface
BALF: Bronchoalveolar lavage fluid
BEGM: Bronchial epithelial growth medium
CFTR: Cystic fibrosis transmembrane regulator
CREC: Conditionally reprogrammed epithelial cell
ELF: Epithelial lining fluid
ELISA: Enzyme linked immunosorbent assay
GGC: γ-glutamylcysteine
GLRX: Glutaredoxin
GPx: Glutathione peroxidase
GSH: Glutathione
HAEC: Human airway epithelial cell
HBEC: Human bronchial epithelial cell
HNEC: Human nasal epithelial cell
HPLC: High performance liquid chromatography
IPA: Ingenuity Pathway Analysis
LC-MS/MS: Liquid chromatography tandem mass spectrometry
LPS: Lipopolysaccharide
NAC: N-acetylcysteine
PIC: Pre-initiation complex
ROS: Reactive oxygen species
SG: Stress granule
SOD: Superoxide dismutase

## Acknowledgements

We thank the study participants and their families for their contributions. We appreciate the assistance from Sydney Children’s Hospitals (SCH) Randwick respiratory department in organization and collection of patient biospecimens – special thanks to Dr John Widger, Dr Laura Fawcett, Dr Yvonne Belessis, Leanne Plush, Amanda Thompson and Rhonda Bell.

## Author contributions

Conceptualization, design and supervision: SAW. Funding acquisition: SAW, AJ, WB. Provision of recourses: AJ, WB, SRT, RMW. Patient samples processing: CKH, NTA. Experimental work: CKH, SLW, BF, LZ. Data analysis and interpretation: CKH, AC, EP, GHS, SAW. Manuscript writing: CKH, AC, SAW with intellectual input from all authors.

---

**Figure S1. Validation of physiological response of HAEC model to LPS challenge. (A)** Primary conditionally reprogrammed epithelial cell lines (CREC) were generated from CF and non-CF controls. **(B)** CREC cells retain their epithelial morphology and express epithelial markers when established with 2D-conventional epithelial culture methods and **(C)** achieve mucociliary differentiation when established at the air liquid interface (ALI). **(D)** Cells exposed to an increasing concentration (10 μg/mL to 100 ug/mL) of LPS show a dose-dependent increase in reactive oxygen species (ROS) production. **(E)** LPS caused significant inflammation via a four-fold increase in pro-inflammatory cytokine IL-8.

**Figure S2. GGC increases cell viability of unchallenged and LPS-challenged immortalised human bronchial epithelial cells.** Cell viability was assessed in (**A**) unchallenged treated, **(B)** post-LPS^+^ treated (therapeutic) and (**C**) pre & post LPS^+^ treated (T+P) conditions in CF immortalised HBEC with DF508/DF508 CFTR genotype (n =3) using the MTT assay. GGC, NAC and GSH treatment in a range of concentration (2 to 2000μl) were tested. DMSO used as positive cell death control. Statistical analysis was performed using one-way ANOVA against the PBS control. (* = p<0.05, ** = p<0.01, *** = p<0.001, **** = p<0.0001). Error bars represent SD.

**Table S1. Antibodies used in study**

**Table S2. Proteomics analysis statistics.**

**Table S3. Differential protein abundance analysis performed with Perseus.**

**Table S4. Canonical pathway, network and disease and biological function analysis perofmed with IPA.**

## References

1. Jones, D.P., et al., Redox state of glutathione in human plasma. Free Radical Biology and Medicine, 2000. 28(4): p. 625–635.

2. Gould, N.S. and B.J. Day, Targeting maladaptive glutathione responses in lung disease. Biochemical pharmacology, 2011. 81(2): p. 187–193.

3. Kogan, I., et al., CFTR directly mediates nucleotide-regulated glutathione flux. The EMBO journal, 2003. 22(9): p. 1981–1989.

4. Dickerhof, N., et al., Oxidative stress in early cystic fibrosis lung disease is exacerbated by airway glutathione deficiency. Free Radical Biology and Medicine, 2017. 113: p. 236–243.

5. Kleme, M.-L. and E. Levy, Cystic fibrosis-related oxidative stress and intestinal lipid disorders. Antioxidants & redox signaling, 2015. 22(7): p. 614–631.

6. Cutting, G.R., Cystic fibrosis genetics: from molecular understanding to clinical application. Nature reviews. Genetics, 2015. 16(1): p. 45–56.

7. Griese, M., et al., Inhalation Treatment with Glutathione in Patients with Cystic Fibrosis. A Randomized Clinical Trial. American Journal of Respiratory and Critical Care Medicine, 2013. 188(1): p. 83–89.

8. Conrad, C., et al., Long-term treatment with oral N-acetylcysteine: Affects lung function but not sputum inflammation in cystic fibrosis subjects. A phase II randomized placebo-controlled trial. Journal of Cystic Fibrosis, 2015. 14(2): p. 219–227.

9. Hayashi, H., et al., Effects of γ-glutamylcysteine ethyl ester on heart mitochondrial creatine kinase activity: involvement of sulfhydryl groups. European Journal of Pharmacology, 1998. 349(1): p. 133–136.

10. Kobayashi, H., et al., The effects of gamma-glutamylcysteine ethyl ester, a prodrug of glutathione, on ischemia-reperfusion-induced liver injury in rats. Transplantation, 1992. 54(3): p. 414–8.

11. Drake, J., et al., Elevation of mitochondrial glutathione by γ-glutamylcysteine ethyl ester protects mitochondria against peroxynitrite-induced oxidative stress. Journal of Neuroscience Research, 2003. 74(6): p. 917–927.

12. Chandler, S.D., et al., Safety assessment of gamma-glutamylcysteine sodium salt. Regul Toxicol Pharmacol, 2012. 64(1): p. 17–25.

13. Zarka, M.H. and W.J. Bridge, Oral administration of γ-glutamylcysteine increases intracellular glutathione levels above homeostasis in a randomised human trial pilot study. Redox Biology, 2017. 11(Supplement C): p. 631–636.

14. Krämer, A., et al., Causal analysis approaches in ingenuity pathway analysis. Bioinformatics, 2014. 30(4): p. 523–530.

15. Yeung, A.W., et al., Role of indoleamine 2, 3-dioxygenase in health and disease. Clinical science, 2015. 129(7): p. 601–672.

16. Chen, L. and B. Liu, Relationships between Stress Granules, Oxidative Stress, and Neurodegenerative Diseases. Oxidative Medicine and Cellular Longevity, 2017. 2017: p. 10.

17. Vidal, L.S., et al., Review of NAD (P) H-dependent oxidoreductases: Properties, engineering and application. Biochimica et Biophysica Acta (BBA)-Proteins and Proteomics, 2018. 1866(2): p. 327–347.

18. Wood, L.G., et al., Oxidative stress in cystic fibrosis: dietary and metabolic factors. Journal of the American College of Nutrition, 2001. 20(2): p. 157–165.

19. Yang, Y., et al., γ-glutamylcysteine exhibits anti-inflammatory effects by increasing cellular glutathione level. Redox biology, 2019. 20: p. 157–166.

20. Le, T.M., et al., γ-Glutamylcysteine ameliorates oxidative injury in neurons and astrocytes in vitro and increases brain glutathione in vivo. Neurotoxicology, 2011. 32(5): p. 518–525.

21. Ferguson, G. and W. Bridge, Glutamate cysteine ligase and the age-related decline in cellular glutathione: the therapeutic potential of γ-glutamylcysteine. Archives of biochemistry and biophysics, 2016. 593: p. 12–23.

22. Quintana-Cabrera, R., et al., γ-Glutamylcysteine detoxifies reactive oxygen species by acting as glutathione peroxidase-1 cofactor. Nature Communications, 2012. 3: p. 718.

23. Nakamura, Y.K., M.A. Dubick, and S.T. Omaye, γ-Glutamylcysteine inhibits oxidative stress in human endothelial cells. Life Sciences, 2012. 90(3): p. 116–121.

24. Braidy, N. and B.-e. Jugder, The precursor to glutathione (GSH),[-glutamylcysteine (GGC), can Ameliorate Oxidative Damage and Neuroinflammation Induced by Amyloid-beta Oligomers in Primary Adult Human Brain Cells. Frontiers in aging neuroscience, 2019. 11: p. 177.

25. Koyama, S., et al., The potential of various lipopolysaccharides to release IL-8 and G-CSF. Am J Physiol Lung Cell Mol Physiol, 2000. 278(4): p. L658–66.

26. Muselet-Charlier, C., et al., Enhanced IL-1β-induced IL-8 production in cystic fibrosis lung epithelial cells is dependent of both mitogen-activated protein kinases and NF-κB signaling. Biochemical and Biophysical Research Communications, 2007. 357(2): p. 402–407.

27. Golebski, K., et al., FcγRIII stimulation breaks the tolerance of human nasal epithelial cells to bacteria through cross-talk with TLR4. Mucosal Immunology, 2019.

28. Aggarwal, S., et al., Glutathione supplementation attenuates lipopolysaccharide-induced mitochondrial dysfunction and apoptosis in a mouse model of acute lung injury. Frontiers in physiology, 2012. 3: p. 161–161.

29. Maiorino, M., M. Conrad, and F. Ursini, GPx4, lipid peroxidation, and cell death: discoveries, rediscoveries, and open issues. Antioxidants & redox signaling, 2018. 29(1): p. 61–74.

30. Yant, L.J., et al., The selenoprotein GPX4 is essential for mouse development and protects from radiation and oxidative damage insults. Free Radical Biology and Medicine, 2003. 34(4): p. 496–502.

31. Castellani, S., et al., NHERF1 and CFTR restore tight junction organisation and function in cystic fibrosis airway epithelial cells: role of ezrin and the RhoA/ROCK pathway. Laboratory investigation, 2012. 92(11): p. 1527–1540.

32. Xie, W., et al., Resolvin D1 reduces deterioration of tight junction proteins by upregulating HO-1 in LPS-induced mice. Laboratory Investigation, 2013. 93(9): p. 991–1000.

33. Kaniuk, N.A., et al., ALIS are Stress-Induced Protein Storage Compartments for Substrates of the Proteasome and Autophagy AU - Szeto, Jason. Autophagy, 2006. 2(3): p. 189–199.

34. Basu, M., S.C. Courtney, and M.A. Brinton, Arsenite-induced stress granule formation is inhibited by elevated levels of reduced glutathione in West Nile virus-infected cells. PLoS pathogens, 2017. 13(2): p. e1006240.

35. Martinovich, K.M., et al., Conditionally reprogrammed primary airway epithelial cells maintain morphology, lineage and disease specific functional characteristics. Scientific Reports, 2017. 7(1): p. 17971.

36. Hamm, J.N., et al., Unexpected host dependency of Antarctic Nanohaloarchaeota. Proceedings of the National Academy of Sciences, 2019. 116(29): p. 14661–14670.

37. Cox, J. and M. Mann, MaxQuant enables high peptide identification rates, individualized p.p.b.-range mass accuracies and proteome-wide protein quantification. Nat Biotechnol, 2008. 26(12): p. 1367–72.

38. Cox, J., et al., Andromeda: a peptide search engine integrated into the MaxQuant environment. J Proteome Res, 2011. 10(4): p. 1794–805.

39. Cox, J., et al., Accurate Proteome-wide Label-free Quantification by Delayed Normalization and Maximal Peptide Ratio Extraction, Termed MaxLFQ. Molecular & Cellular Proteomics, 2014. 13(9): p. 2513–2526.

40. Tyanova, S., et al., The Perseus computational platform for comprehensive analysis of (prote) omics data. Nature methods, 2016. 13(9): p. 731.

