## Supplementary figures and images for "Novel antioxidant therapy with the immediate precursor to glutathione, γ-glutamylcysteine (GGC), ameliorates LPS-induced cellular stress in an *in vitro* cystic fibrosis model"

### Figure S1

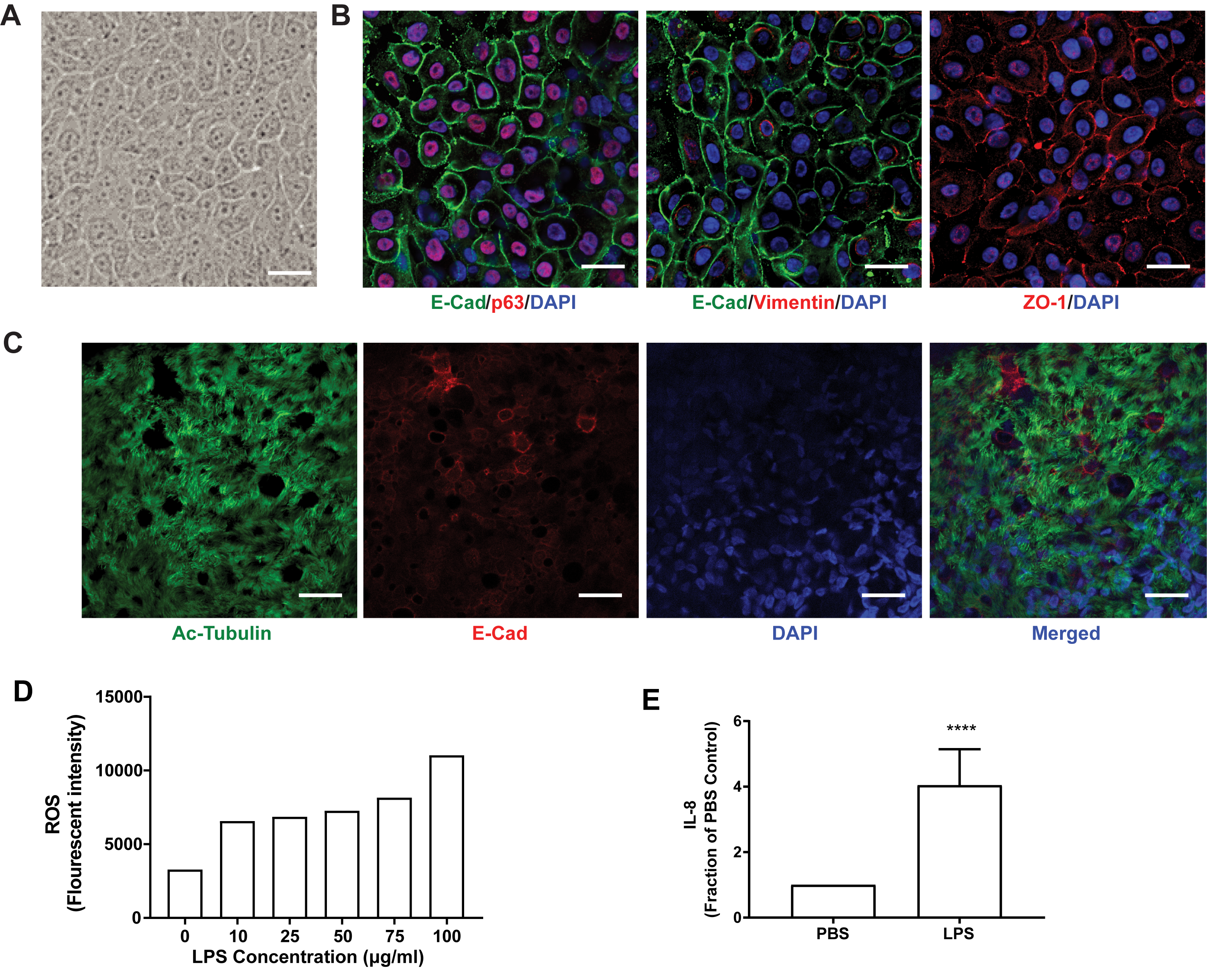

### Figure S2

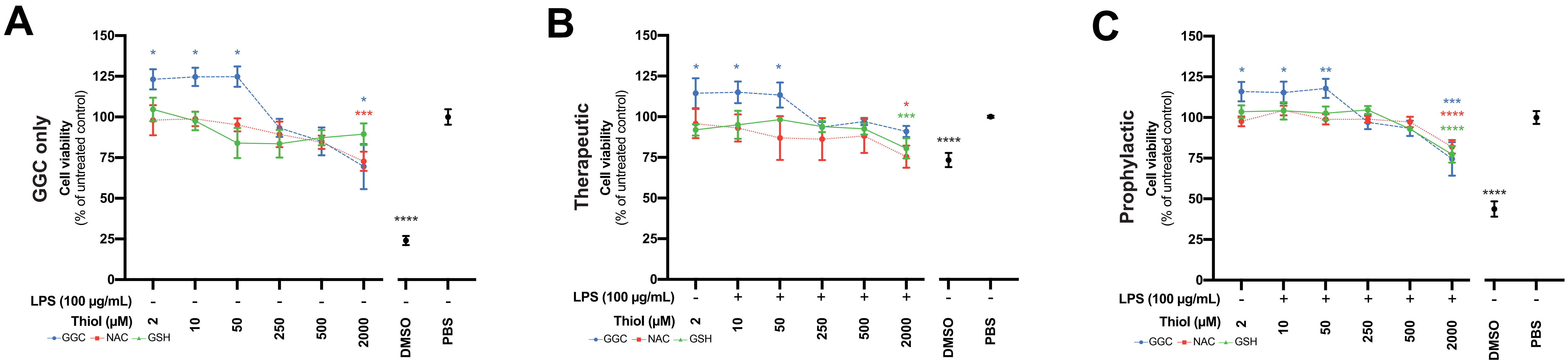
