## Supplementary material for "Novel antioxidant therapy with the immediate precursor to glutathione, γ-glutamylcysteine (GGC), ameliorates LPS-induced cellular stress in an *in vitro* cystic fibrosis model": Table S1

**Table S1. Antibodies used in study**

| **Antibody** | **Supplier** | **Dilution** | **Marker** | **Experiment** |
| --- | --- | --- | --- | --- |
| Epithelial-Cadherin (E-cad) | Invitrogen (13-1700) | 1:100 | Epithelial cell adhesion protein | Epithelial cell line validation |
| Vimentin | Abcam (ab92547) | 1:250 | Intermediate filament protein expressed in mesenchymal cells | Epithelial cell line validation |
| ZO-1 | Invitrogen (61-7300) | 1:50 | Tight junction protein | Epithelial cell line validation & ZO-1 immunofluorescence |
| p63 | Abcam (ab124762) | 1:250 | Basal cell marker | Epithelial cell line validation |
| Epithelial-Cadherin (E-cad) | Cell Signaling (3195) | 1:100 | Epithelial cell adhesion protein | ALI culture validation |
| Acetylated tubulin | Sigma  (T7451) | 1:250 | Marker for cilia | ALI culture validation |
| EIF4A1 | Abcam (ab31217) | 1:1000 | Stress granule protein | Stress granule immunofluorescence |
| G3BP | Abcam (ab56574) | 1:100 | Stress granule protein | Stress granule immunofluorescence |
| Alexa Fluor 488 goat anti-mouse IgG (H+L) | Invitrogen (A-11029) | 1:500 | Secondary antibody |  |
| Alexa Fluor 647 goat anti-rabbit IgG (H+L) | Invitrogen (A-21245) | 1:500 | Secondary antibody |  |
| Alexa Fluor anti-mouse 555 | Invitrogen (A-21424) | 1:500 | Secondary antibody |  |
