## Supplementary material for "Novel antioxidant therapy with the immediate precursor to glutathione, γ-glutamylcysteine (GGC), ameliorates LPS-induced cellular stress in an *in vitro* cystic fibrosis model": Table S2

**Table S2. Proteomics analysis statistics.**

| **Comparison (condition 1 vs. condition 2)** | **Number of proteins (cond. 1)** | **Number of proteins (cond. 2)** | **Total Shared Proteins** | **Significantly Changed Proteins** | **Significantly Up Proteins (Cond. 1 relative to Cond. 2)** | **Significantly Down Proteins (Cond. 1 relative to Cond. 2)** |
| --- | --- | --- | --- | --- | --- | --- |
| Mock vs. LPS^+^ | 1858 | 1836 | 1493 | 65 | 44 | 21 |
| LPS^+^ vs. Therapeutic | 1836 | 1908 | 1526 | 56 | 18 | 38 |
| LPS^+^ vs. Prophylactic | 1836 | 1951 | 1561 | 57 | 28 | 29 |
| LPS^+^ vs. T+P | 1836 | 1719 | 1402 | 39 | 15 | 24 |
| Mock vs. LPS^-^ Tx | 1858 | 1884 | 1481 | 37 | 26 | 11 |
